# Draft genome of *Palmaria palmata* and intraspecific genetic variations in the North-East Atlantic

**DOI:** 10.1101/2024.07.05.602248

**Authors:** Serena Rosignoli, Masami Inaba, Matthias Schmid, Marcus McHale, Antoine Fort, Maeve D. Edwards, Agnes Mols Mortensen, Roy Bartle, Arild Endal, Aurélien Baud, Christine Maggs, Ronan Sulpice

## Abstract

The rhodophyte *Palmaria palmata* (L.) Weber & Mohr is one of the target species of a growing European seaweed industry due to its high content of protein and essential amino acids which makes it suitable for human food, dietary supplements, and as salmon feed. However, the lack of a published nuclear genome limits phylogenetics analyses and gene function investigations which could help the development of a breeding programme.

We present the first draft genome of *P. palmata* that was obtained with PacBio HiFi long read sequencing with average coverage of 10×, consisting of an assembly of 1.05 Gb, N50=2.75Mb and BUSCO completeness of 72.1%. Additionally, a population study on the whole genome of 33 *P. palmata* individuals from across the Northern East Atlantic area found three main clusters consistent with their geographic distribution: (1) Denmark and Norway, (2) France and western Ireland, (3) Faroe Islands. All individuals from Northern Ireland share ancestry with western Ireland and Denmark, and some individuals from the Faroe Islands show admixture from Faroe, western Ireland and Northern Ireland. These results represent a fundamental step towards breeding and genetic studies to further explore the vastly unexploited economic potential of *Palmaria palmata*.

**Highlights:**

- We report the first draft genome of *Palmaria palmata*, from PacBio HiFi long reads.
- The size of the genome, 1.05 Gb, is among the largest so far among the Rhodophyta.
- Busco completeness of 72.1% and contig N50 of 2.75 Mb indicate good quality.
- The genomes of 33 more individuals from North Atlantic Europe have been sequenced.
- Phylogenetic analysis found three clusters consistent with geographic distribution.

## Introduction

The red seaweed *Palmaria palmata*, commonly known as dulse, has significant economic and ecological value and is one of the target species of a growing European seaweed aquaculture industry [1]. This species is widely distributed across the North Atlantic, from its southern distribution limits in Portugal [2] and Long Island (40°N)[3] up to polar regions in Svalbard [4] and the north of Greenland (80°N) (GBIF.Org 2024). *Palmaria palmata* is one of the few seaweed species that has been traditionally consumed in western countries [1], [6], [7], [8], and more recently it has also been used as food supplement and salmon feed. It has a rich nutritional profile, including high levels of proteins [9], [10] essential amino acids [11] and a fatty acid profile rich in omega-3 fatty acids [12], [13]. It has an unusual heteromorphic diplohaplontic life cycle that complicates its cultivation. The tetrasporophyte grows up to 10-20 cm and is morphologically similar to the male gametophyte, it can only be distinguished when it develops sori. The female gametophyte measures only a few millimetres, is crustose and become sexually mature a few days after tetraspore release, while the male gametophyte requires 9-12 months before forming spermatia, so females are fecundated by males from the previous gametangial generations. The tetrasporophyte develops on the fertilized female and overgrows it [14]. The majority of commercially used *P. palmata* comes from harvesting wild populations [15]. However, due to the demand for high quality *P. palmata* biomass, there is also an increased focus on commercial cultivation of this species, for example in Ireland, Denmark and France [16], [17]. This species can be grown in onshore tanks or on rope cultures offshore (Stévant et al. 2023). Preliminary studies have shown variability in traits like growth rate and protein content and have demonstrated the potential of strain selection (Matthias Schmid, personal communication). Despite its commercial importance, no nuclear reference genome is available for *P. palmata* yet, which hinders breeding efforts. Previous studies have shown that genome size in other Rhodophyta varies between ca. 9-1,460 Mb. The number of protein coding genes is between ca. 5,000 and 13,000, the vast majority being single exon genes, much lower than in land plants, which usually contain over 30,000 coding genes [19], [20], [21], [22]. It is assumed that Rhodophyta went through a genome reduction caused by an adaptation to extreme environments and a subsequent genome expansion by transposable elements [23], [24], [25].

Genetic and phylogenetic studies in *P. palmata* are limited. So far, only one nuclear gene has been cloned in *P. palmata* [26]. Using a combination of nuclear, plastid and mitochondrial genetic markers Provan, Remi and Maggs [3] studied the biogeographic history of *P. palmata* in the North Atlantic, which revealed the existence of a previously unidentified marine refugium in the English Channel, along with possible secondary refugia off the southwest coast of Ireland and in northeast North America and/or Iceland. Such phylogenetic insights can illuminate patterns of genetic variation and adaptation across different geographic regions, which is essential for both conservation efforts and the strategic development of region-specific cultivation practices [27], [28], [29].

The main aim of this study was to obtain the first draft genome of *P. palmata* using PacBio HiFi sequencing. Moreover, to test the genome and assess genetic variability in the North-East Atlantic, we conducted a phylogenetic analysis based on the genomes of 33 tetrasporophytes sequenced with Illumina. The draft genome of *P. palmata* marks a pivotal step in unlocking the genetic potential of this commercially valuable red seaweed. The integration of genomic data with phylogenetic analysis from multiple geographic regions offers an understanding of the species’ genetic diversity and evolutionary adaptation. These insights pave the way for improvements for the cultivation and further commercialization of *P. palmata*, aligning with the growing demand for sustainably sourced and high-quality natural products.

## Materials and methods

### Samples

For the genome assembly, a male gametophyte was obtained from tetraspores of *Palmaria palmata* originally collected at Castlegregory, Kerry, Ireland (52.260889, -10.013500) on 14/12/2020. Tetraspores were settled and grown on culture string and held in stasis conditions within the laboratory (half-strength F/2 culture medium held at 10 °C, light intensity and photoperiod: 20 µmol m^-2^ sec^-1^, 12:12 light:dark) until 8/2/2022 when the cultures of male gametophytes were deployed at sea on longlines at the Marine Institute Lehenagh Pool Marine Research Site, Co. Galway (53.400720, -9.819379). Samples of whole fronds were harvested 8/4/2022 and returned to laboratory conditions to grow for a further month to obtain an appropriate biomass for sampling (full-strength F/2 culture medium held at 10 °C, light intensity and photoperiod: 90 µmol m^-2^ sec^-1^, 16:8 light:dark).

For the phylogenetic study, 33 tetrasporophytes of *Palmaria palmata* were collected from 10 locations in 6 different countries in Europe: Norway, Denmark, Faroe Islands, Northern Ireland, Ireland and France (Table 1). They were all wild harvested except the samples from Denmark, cultivated on land by Pure Algae, Grenaa, Denmark.

**Table 1.** Tetrasporophyte samples used in the phylogenetic study.

| Location | Code | Latitude | Longitude | Number of individuals | Collector/sender | Date of collection | Individuals ID |
| --- | --- | --- | --- | --- | --- | --- | --- |
| Aursvika, Malangen, Norway | NOR.1 | 69.313680 | 18.665561 | 1 | Arild Endal, Tromsø, Norway | 25/04/2021 | P44 |
| Loddbukta, Balsfjord, Norway | NOR.2 | 69.233512 | 19.374789 | 3 | Arild Endal, Tromsø, Norway | 25/04/2021 | P39, P41, P42 |
| Fornæs, Grenå, Denmark | DNK.1 | 56.442472 | 10.954555 | 3 | Lasse Hornbek Nielsen, Pure Algae Denmark | 20/02/2021 | P33, P36, P38 |
| Fámjin, Suðuroy Faroe Islands | FRO.1 | 61.526944 | -6.881667 | 4 | Agnes Mols Mortensen, TARI Faroe Seaweed, Faroe Islands | 04/09/2020 | P6, P7, P12, P13 |
| Hvítanes, Streymoy, Faroe Islands | FRO.2 | 62.043333 | -6.774722 | 4 | Agnes Mols Mortensen, TARI Faroe Seaweed, Faroe Islands | 04/09/2020 | P2, P3, P8, P9 |
| Oyrargjógv, Faroe Islands | FRO.3 | 62.105833 | -7.149444 | 4 | Agnes Mols Mortensen, TARI Faroe Seaweed, Faroe Islands | 04/09/2020 | P4, P5, P10, P11 |
| Ballycastle, Co. Antrim, Northern Ireland | NIR.1 | 55.211328 | -6.195065 | 3 | Ruairidh Morrison, North Coast Smokehouse, Ballycastle, UK | 26/02/2021 | P32 |
| Ballyvaughan, Co. Clare, Ireland | IRL.2 | 53.118941 | -9.15141 | 1 | Antoine Fort, Technological University of the Shannon, Ireland | 14/07/2020 | P1 |
| Roundstone, Co. Galway, Ireland | IRL.1 | 53.397343 | -9.91610 | 7 | Antoine Fort, Technological University of the Shannon, Ireland | 05/10/2020 | P14-P20 |
| Saint-Brieuc, France | FRA.1 | 48.524389 | -2.713556 | 3 | Aurélien Baud, Station Biologique de Roscoff, France | 25/02/2021 | P21-P23 |

### Genome Assembly

For the assembly, total DNA of the male gametophyte was extracted with Qiagen Genomic-Tips (Qiagen, Courtaboeuf, France) and sequenced on the PacBio Sequel 2 platform (Pacific Biosciences Inc., CA, USA) with circular consensus sequencing [30] by Novogene Co., Ltd. (UK). To estimate genome size and ploidy level, Kmer analysis was performed with KAT v.2.4.2 [31]. HiFi reads were assembled with hifiasm [32]. Completeness was assessed with BUSCO v. 5.4.4 [33], using the eukaryota_odb10 datasets with 255 markers. GC content was evaluated with the kat command *sect* and coverage with bedtools v.2.27.1 [34], command *genomecov*. For contamination analysis, assembly regions were assigned a taxonomy classification by Kraken 2 [35], using the database PlusPFP version 09/12/2022; contigs classified as prokaryotic were removed with seqtk v1.3 [36]. Repetitive DNA elements were identified with RepeatModeler2 [37] and soft masked with RepeatMasker. Gene prediction was carried out with a combined approach through BRAKER3 [38], integrating homology based prediction and RNA-Seq data of *P. palmata* publicly available in Sequence Read Archive (SRA, https://www.ncbi.nlm.nih.gov/sra/) accession SRX2867140, downloaded on 12/01/2024. RNA-Seq reads were aligned to the masked assembly genome with STAR v.2.7.8 [39]. Predicted sequences were blasted against the UniProt/SwissProt database, release 2023_02 (The UniProt Consortium 2023) and against the 82,161 protein sequences of Florideophyceae from NCBI RefSeq [41], downloaded on 18/03/2023. Functional annotation of protein-coding genes was obtained with InterProScan 5 v.5.56.89 [42]. The final GTF file was composed with *agat_sp_manage_functional_annotation*.*pl* and filtered with agat_sp_keep_longest_isoform.pl [43].

### Phylogenetic analysis

DNA of the 33 tetrasporophytes was extracted with either the SILEX [44] or the magnetic bead protocol [45] and sequenced with Illumina Novaseq PE 150 by Novogene (UK). For the phylogenetic analysis, Illumina reads of the 33 individuals were aligned to our new draft genome with BWA v. 7.17 command *mem*, and Samtools v. 1.13 command *view* [46], [47]. Variants were called with bcftools v. 1.5 [48] and filtered for: minimum base quality 20, minimum mapping quality 30, minimum phred-scaled probability QUAL 40 and depth between 10 and 70. For the organellar genomes, reads were aligned to the available references [49], [50] and variant calling was performed as described above, except minimum depth was set to 200×. The SNP matrix for the nuclear genome was obtained with bcftools *merge* and filtered for minimum allele count 3 and maximum missing rate of 0.5. Discriminant Analysis of Principal components (DAPC) was conducted on the SNP matrix using 4 principal components and 4 discriminate analyses. The results were visualized with RStudio v. 2022.12.0 and the R package *adegenet* v.2.1.10 [51].

Population structure in the set of 33 individuals was evaluated with Admixture v.1.3 [52] and visualized with Pong [53]; principal component analysis was run with EIGENSOFT smartpca v.8 [54], [55] and visualized with RStudio. Discriminant analysis of principal components (DAPC) was performed using the R package *adegenet* v2.1.10 [51] for 1 to 33 clusters, and the Bayesian information criterion (BIC) was used to identify the optimal number of clusters to describe the population. Maximum likelihood phylogenetic study was inferred with RAxML-NG v. 1.2.0 [56] with bootstrap resampling and substitution model GTR+G; the vcf file containing the SNP matrix was converted into phylip format with the script vcf2phylip v. 2.0 [57]; bootstrap convergence was verified post-hoc with cutoff values from 0.01 to 0.05.

## Results

### Assembly of a male gametophyte reveals large genome size compared to other Rhodophyta

PacBio circular consensus sequencing of the male gametophyte produced 614,981 HiFi reads, for a total length of 12.9 Gb, and a read N50 of 21.5 Kb. Haploidy was confirmed by k-mer analysis (k=21) which found 9,472,219,659 k-mers and estimated an average k-mer coverage of 9× and a genome size of ∼ 1.17 Gb (Suppl. Fig. S1). The assembly generated with hifiasm consisted of 1,469 contigs, with a total length of 1,122,823,132 bp, average coverage 10×, contig N50 2Mb, and the largest contig 13.81 Mb. The BUSCO completeness was 75.7% and the missing rate 15.7% (n=255). The average genome coverage calculated with bedtools was 10.7×. The GC content was 51.7% and 6.5% of the genome size was likely of prokaryotic origin. Hence, the corresponding 454 contigs, total length 72,515,266 bp, were discarded. The most abundant contaminant bacterial phyla (Suppl. Fig. S2) were Pseudomonadota (42.6% of total contaminants length), Actinomycetota (32.2%), Cyanobacteriota (12.0%), Bacteroidota (6.0%), Mixococcota (2.8%) and Bacillota (2.2%).

After removal of prokaryotic contigs, the resulting DNA assembly consisted of 1015 contigs, with a total length of 1,050,307,866 bp, a BUSCO completeness of 72.1% and a missing rate 20.4% (n=255). N50 was 2.75 Mb and the GC content 51.9% (Table 2). The Repetitive elements constituted 91.3% of the whole genome size (Suppl. Table S1), of which 90.89% were interspersed repeats. Retrotransposons, in particular LTR elements, were the most abundant (44.27%), the DNA transposons constituted 23.85% of the genome and 22.77% were unclassified. 9,641 protein coding genes, with an average gene length of 2,376 bp, were annotated with a combined approach. The total gene length was 68,562,153 bp, equivalent to 6.5% of the entire genome. A high proportion of protein-coding genes (66%) were single exon genes, 91.6% of protein coding genes had at least one INTERPRO associated and 72% of them had a gene ontology (GO) annotation. In comparison with other published genomes of red seaweeds (Suppl. Table S2), a N50 of 2.75 Mb is among the highest in the Rhodophyta, exceeded only by *Pyropia haitanensis*, indicating a good contiguity of the assembly. At 1.05 Gb, the genome of *P. palmata* is the third largest known in the Rhodophyta.

**Table 2.** Summary statistics of the *Palmaria palmata* genome.

| **HiFi reads** |  |
| --- | --- |
| Total length of reads (bp) | 12,862,012,646 |
| Nr. of HiFi reads | 614,981 |
| Average read length (bp) | 20,914 |
| N50 reads (Kb) | 21.5 |
| Average coverage (×) | 10 |
| **Genome assembly** |  |
| Genome size (bp) | 1,050,307,866 |
| **Contigs** |  |
| Nr. of contigs | 1,015 |
| Largest contig size (Mb) | 13.81 |
| N50 (Mb) | 2.75 |
| L50 | 120 |
| GC content (%) | 51.86 |
| **BUSCO evaluation of assembly** |  |
| Completeness, n=255 | C:72.1%[S:63.1%,D:9.0%],<br>F:7.5%,M:20.4% |
| **Gena annotation** |  |
| Number of protein-coding genes | 9,641 |

### Phylogenetic analyses of 33 individuals

#### Variant calling on 33 whole genome sequences reveal variation across Northern Europe

The alignment of Illumina reads and subsequent variant calling of the 33 tetrasporophyte genomes compared to our new draft genome produced a SNP matrix of 9,885 SNPs (Suppl. File) with minimum allele count 3 and maximum missing rate of 0.5. The number of SNPs per individual ranges from 924 for P1 from Ballyvaughan, Ireland, to 4,944 for P38 from Fornæs, Denmark (Suppl. Table S3). 34 SNPs appear to be unique for a single location: 1 for Hvítanes, Faroe Islands, 16 for Fornæs, Denmark, 3 for Ballycastle, Northern Ireland, 14 for Saint-Brieuc, France (Suppl. Table S4). No genetic variability was found among our individuals for the gene *ppKab*, the only protein coding nuclear gene cloned in *P. palmata* so far, which is involved in kainic acid biosynthesis [26].

### Population structure shows three main clusters

Admixture proportion inference was evaluated for a number of ancestral populations (k) from 1 to 33, where 33 is the number of individuals (Fig. 1.a). An optimal number of population clusters of 3 was estimated with the Bayesian Information Criterion (BIC), (Suppl. Fig. S3). Considering 3 ancestral populations, all individuals from Denmark and Norway (n=7) descend from the same population *Pop1* (Fig. 1.a, b), all individuals from France and western Ireland (n=11) descend from *Pop2*, 9 individuals out of 12 from Faroe descend from *Pop3*, while 3 individuals from Oyragjógv, Faroe, are admixed and descend mainly from *Pop3* and are also partially assigned to *Pop1* and *Pop2*; all the 3 individuals from Northern Ireland are admixed and descend from both *Pop1* and *Pop3*. DAPC on the SNP matrix for the whole genome of 33 individuals with locations as prior (Fig. 1.c) found genetic similarity within the samples from France and western Ireland, within the Faroe samples, and within samples from both Norway sites, while the individuals from Denmark and Northern Ireland are intermediate between Norway and western Ireland. Both Admixture and DAPC are concordant in clustering individuals from Norway, from France/western Ireland, and the majority of individuals from Faroe Islands. Denmark grouped with Norway from Admixture analysis but was more intermediate in DAPC analysis. In both analyses, Northern Ireland is intermediate between France/western Ireland and Norway.

**Figure 1.**
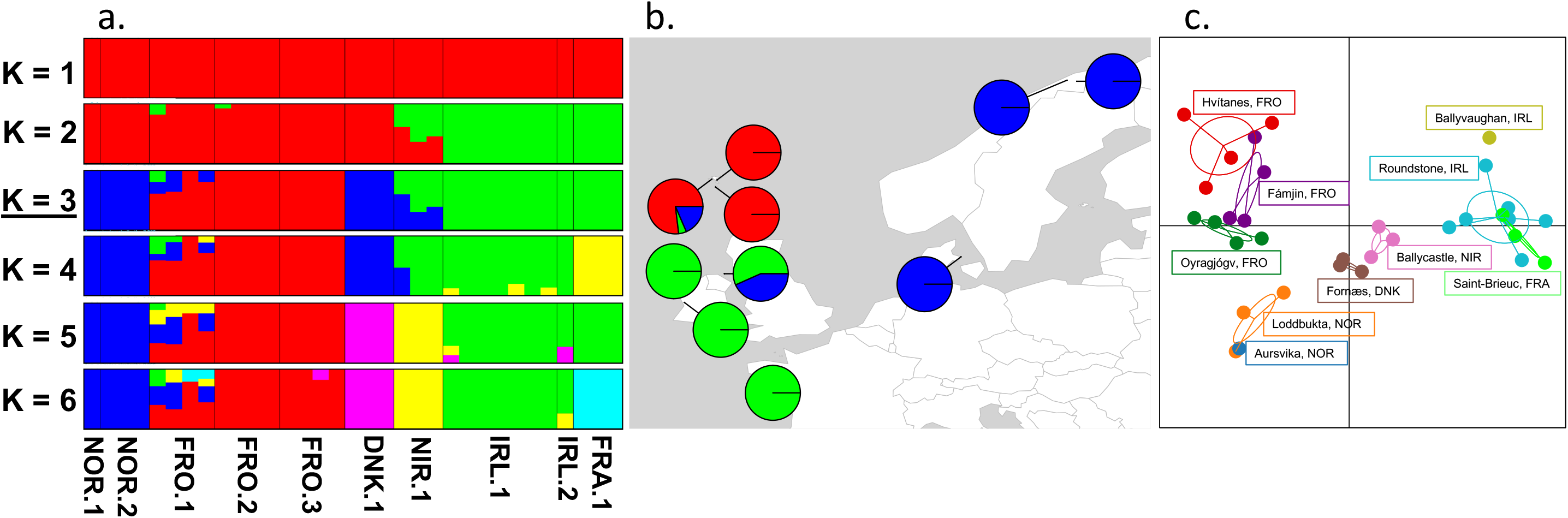
Whole genome population structure of 33 individuals. a. Admixture proportion for ancestry coefficients k from 1 to 6. Every column represents one individual; on the bottom, their geographic provenance is indicated (see Table 1). b. Admixture proportion of each subpopulation for k = 3 on the map of Northern Europe. d. Discriminant Analysis of Principal components (DAPC) using 4 principal components and 4 discriminate analyses.

### Phylogenetic tree

The structure of the maximum likelihood phylogenetic tree (Fig. 2) reflected in large part the geographic origin of individuals: the main (oldest) differentiation was between Norway, Denmark and the rest of the countries involved, thus Norway and Denmark are sister populations. Ireland (west coast) and France are sister populations, and their nearest relative is the Northern Ireland population. Individuals from Faroe might belong to two different lineages, although the support values are lower than 90: the majority of them, 8 out of 10, are in a sublineage of their own and 2 are more closely related to samples from western Ireland, France and Northern Ireland. Overall, the tree suggests a geographical progression of population differentiation from North East to South West. The populations of Norway and Denmark are the most distinct, followed by the populations of the Faroe Islands and Northern Ireland, while the populations of western Ireland and France are genetically closer.

**Figure 2.**
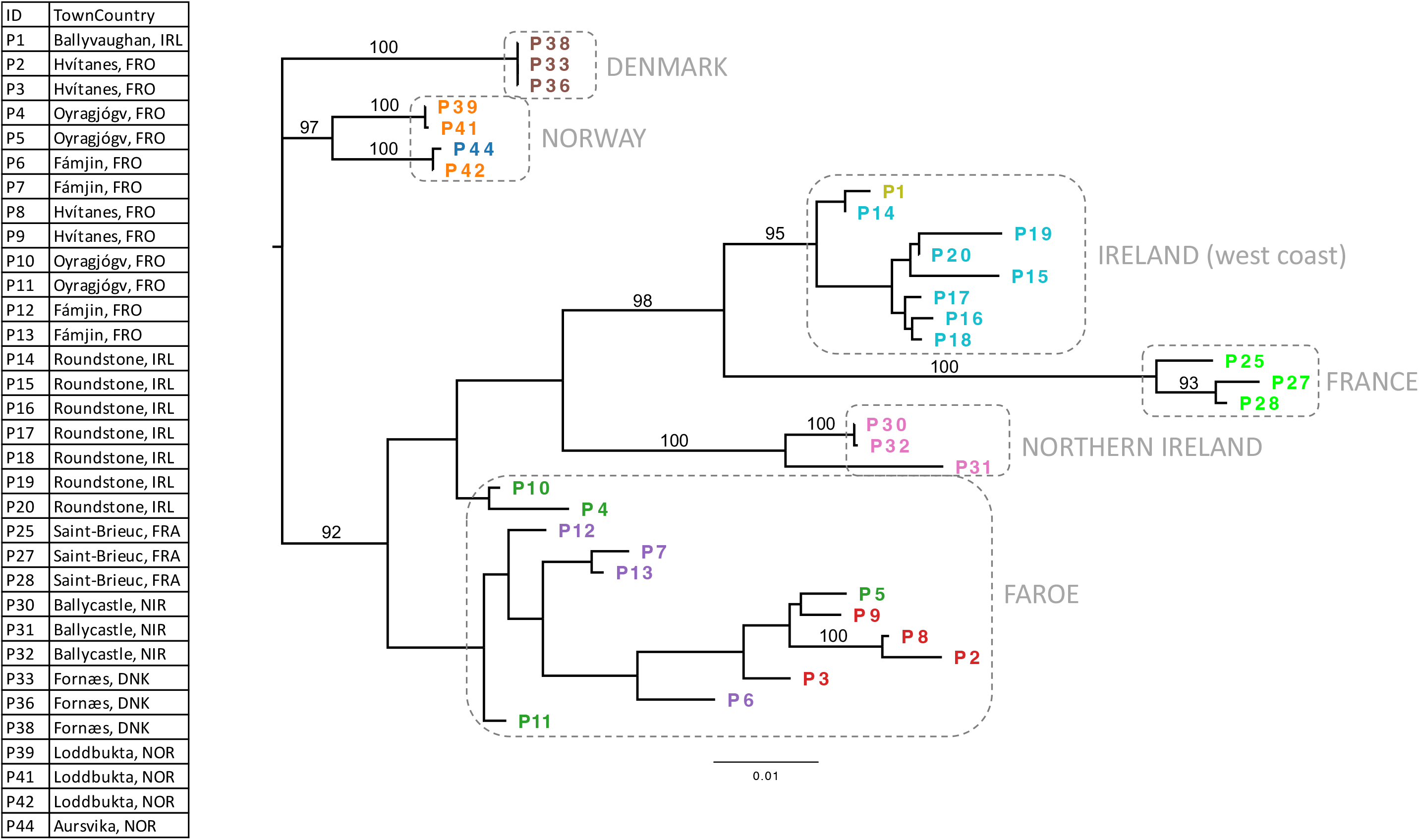
Molecular phylogenetic analysis with the maximum likelihood method on the SNPs of 33 individuals of *P. palmata* with annotation of their geographic origin and colors corresponding to site. On the edges, support values greater than 90 are reported.

## Discussion

### Genome parameters

At 1.05 Gb, the genome of *P. palmata* is among the largest so far out of the 84 available Rhodophyta genomes, preceded only by Bostrychia radicans and Bostrychia flagellifera [58] (Suppl. Table S2) and it is around 36 times the size of the shortest genome among the Florideophyceae, that of *Opuntiella californica*, while Bangiophyceae span from the 8.79 Mb genome of *Cyanidium caldarium* [59] to the 108Mb genome of *Pyropia yezoensis* [60]. The large genome size of *P. palmata* is caused by the high number of transposable elements (TE), in particular of LTR, which is the main determinant of genome size variation in land plants besides polyploidization, as reviewed in [61], and in red algae [23], [24]. In fact, *P. palmata* has the highest percentage of repeat elements (RE) among the sequenced Rhodophyta, 91.3%. Of this, 44.3% of the genome size is retroelements, of which 36.2% are LTR, and 23.9% are DNA transposons. This is twice the number of RE in *Chondria armata* (45.5%), which has a genome that is roughly half the size of *P. palmata*. The GC content of 51.9% falls within the range of other Florideophyceae (45.0-52.5%). The BUSCO score of 72.1% is in the expected range, given the lack of lineage-specific databases for Rhodophyta [20], [62].

### Gene annotation

The number of protein coding genes in *P. palmata*, 9,641, is similar to other Florideophyceae (7,174-11,437) and lower than in land plants (reviewed in Panchy, Lehti-Shiu and Shiu, 2016), in concordance with the hypothesis of gene reduction in the Rhodophyta [25], [64] followed by genome expansion by transposable elements in the Florideophyceae [65]. Single exon genes make up a high proportion of protein-coding genes, 66%, comparable to other Rhodophyta [66].

### Microbial contaminants

The most abundant bacterial phyla before decontamination (Suppl. Fig. S2) were Pseudomonadota (42.6% of total contaminants length), Actinomycetota (32.2%), Cyanobacteriota (12.0%), Bacteroidota (6.0%), Mixococcota (2.8%) and Bacillota (2.2%). This is largely consistent with other studies in Rhodophyta [67], [68], [69], [70], [71], except for Actinomycetota, which were much more represented in *P. palmata* compared to other species, in which they never exceeded 10% [67], [68], [69], [70], [71]. The most abundant genus was *Streptomyces* (21.2% of total contaminants length) followed by *Aeromonas* (7.7%) and *Dickeya* (7.2%). Actinomyces are a rich source of bioactive secondary metabolites. In particular, members of the genus *Streptomyces* from marine environments can have antimicrobial, antifungal and nematicidal activity, and can produce phytohormones like gibberellic acid, indole acetic acid, abscisic acid, kinetin and benzyladenine [72], [73], [74]. *Aeromonas* are ubiquitous aquatic bacteria sometimes associated with foodborne diseases [75], *Dickeya* are mainly known as plant pathogenic bacteria [76].

### Variant calling

The 34 SNPs that are unique for a region or country of origin, can be used as described in [77] for fast and efficient traceability of end products derived from *P. palmata*.

### Population structure and phylogenetic study

Population structure analyses with Admixture and DAPC showed consistent results. In both analyses, individuals from the same location were similar and genetic divergence increased with geographic distance, as was expected (Fig. 1.c). Three main clusters were identified by both methods with minimal differences regarding two locations, Northern Ireland and Denmark, located at the centre of the whole geographic distribution of the sampling. The same clustering is largely consistent with the phylogenetic tree, with the exception of the Faroe Islands which were more closely related to Norway in the Admixture and in the DAPC compared to the phylogenetic tree.

These results are congruent with previous phylogeographic analyses of *P. palmata* [3]. In that work, the authors studied, among other loci, the distribution of plastid haplotypes for two regions, including the genes 5S, 16S and 23S rDNA, rbcL, rbcS, trnI and trnA, and found a similar east-west division in northern Europe: Norway, Denmark with one haplotype, western Ireland and France with another. In their study, Northern Ireland clustered with Norway and Denmark, and Faroe clustered with Ireland and France whilst in our analysis Northern Ireland was a mixture of both groups and Faroe shared a partial contribution from both groups. Furthermore, the plastid rpl12-rps31-rpl9 region of all the individuals in our study was identical corresponding to the haplotype AY826508.1 from GenBank, which in [3] was called P-I, and was found in individuals from the temperate northeast Atlantic, Irish Sea, Bay of Biscay and North America.

## Conclusions

In this work, we obtained the first draft genome of *Palmaria palmata*, a rhodophyte with varied applications and vastly unexploited economic potential. *Palmaria palmata* has the largest genome sequenced in the Rhodophyta so far. The population study highlighted three main genetic clusters within Northern Europe. This work is a fundamental step towards breeding and further genetic studies.

## Credit

Serena Rosignoli: conceptualization, data curation, formal analysis, software, validation, visualization, writing; Masami Inaba: investigation, resources, validation; Matthias Schmid: funding acquisition, writing, validation; Marcus McHale: resources, validation: Antoine Fort: investigation, resources, validation; Maeve D. Edwards: resources, methodology; Agnes Mols Mortensen: resources; Roy Bartle: resources; Arild Endal: resources; Aurélien Baud: resources; Christine Maggs: validation, writing-editing; Ronan Sulpice: conceptualization, funding acquisition, supervision, validation, resources, writing-editing.

## Data availability

The genome has been deposited at DDBJ/ENA/GenBank under the accession JBEXAG000000000.

## Acknowledgements

The authors thank Ruairidh Morrison, Lasse Hornbek Nielsen, Nikoline Ziemer and Jonas Steenholdt Sørensen for the provision of samples of *P. palmata*.

## ABBREVIATIONS

DNK: Denmark
FRA: France
FRO: Faroe Islands
IRL: western Ireland
NIR: Northern Ireland
NOR: Norway.

