## Supplementary for "Draft genome of *Palmaria palmata* and intraspecific genetic variations in the North-East Atlantic"

### Supplementary materials

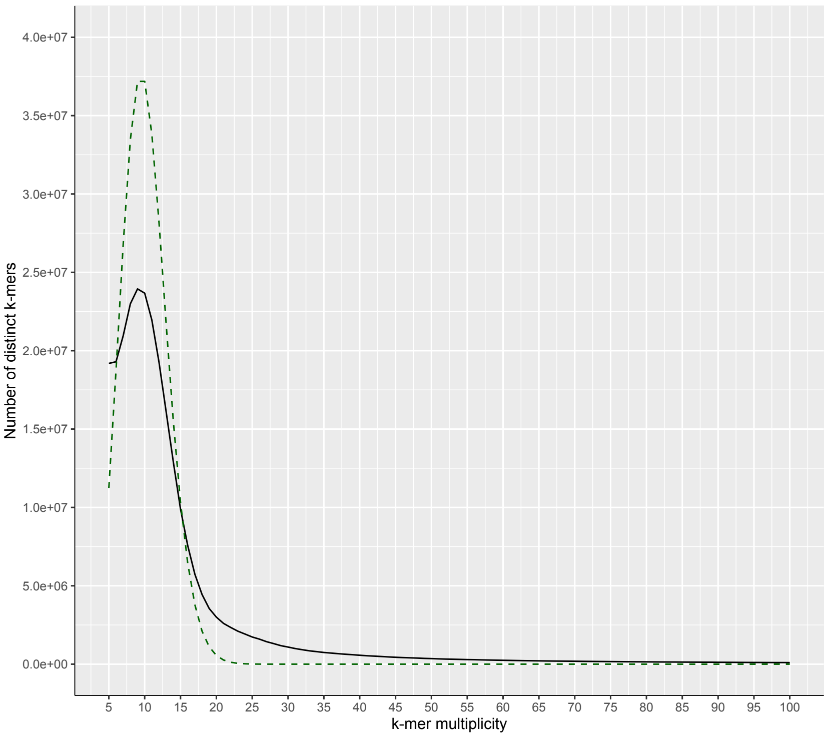


**Supplementary Figure S1. 21-mer spectrum for raw reads of *P. palmata***. The graph shows the number of distinct k-mers as a function of k-mer multiplicity, or frequency. With the choice of k=21, it can be assumed that k-mers are unique in the genome. Values of x lower than 5 correspond to erroneous reads and have been eliminated. The green dashed curve indicates the ideal Poisson distribution. The k-mer spectra curve is regularly shaped, it has one peak, indicating a haploid genome, with maximum at x=9, that is the estimated mean coverage of our genome. The area under the curve represents the total number of k-mers (10,563,588,785); the size of the genome is estimated as: area under the curve/average coverage=1.17 Gb.

**Supplementary Table S1. Repetitive elements in the assembly of *P. palmata*.** The first column lists the type of the repetitive element (RE), the second column the number of the RE in the genome, the third column the total size in bp, and the last column contains the portion of the RE in percentage on the total genome size. RE constitute 91.3% of the genome.

| | | | |
|---|---|---|---|
| **Retroelements** | **403,616** | **464,941,371 bp** | **44.27%** |
| SINEs: | 0 | 0 bp | 0.00 % |
| Penelope | 535 | 75,285 bp | 0.01 % |
| LINEs: | 55,183 | 84,646,902 bp | 8.06 % |
| CRE/SLACS | 25,170 | 27,111,188 bp | 2.58 % |
| L2/CR1/Rex | 696 | 642,953 bp | 0.06 % |
| R1/LOA/Jockey | 0 | 0 bp | 0.00 % |
| R2/R4/NeSL | 97 | 33,244 bp | 0.00 % |
| RTE/Bov-B | 6,713 | 10,515,896 bp | 1.00 % |
| L1/CIN4 | 3,658 | 7,696,336 bp | 0.73 % |
| LTR elements: | 348,433 | 380,294,469 bp | 36.21 % |
| BEL/Pao | 0 | 0 bp | 0.00 % |
| Ty1/Copia | 92,240 | 138,802,962 bp | 13.22 % |
| Gypsy/DIRS1 | 88,675 | 130,509,039 bp | 12.43 % |
| Retroviral | 589 | 244,763 bp | 0.02 % |
| **DNA transposons** | **148,243** | **250,547,870 bp** | **23.85 %** |
| hobo-Activator | 3,644 | 2,999,063 bp | 0.29 % |
| Tc1-IS630-Pogo | 2,288 | 1,023,145 bp | 0.10 % |
| En-Spm | 0 | 0 bp | 0.00 % |
| MULE-MuDR | 55,115 | 120,828,859 bp | 11.50 % |
| PiggyBac | 0 | 0 bp | 0.00 % |
| Tourist/Harbinger | 2,308 | 2,568,609 bp | 0.24 % |
| Other | 0 | 0 bp | 0.00 % |
| Rolling-circles | 267 | 430,935 bp | 0.04 % |
| Unclassified: | 450,557 | 239,181,009 bp | 22.77 % |
| **Total interspersed repeats:** | | **954,670,250 bp** | **90.89 %** |
| Small RNA: | 292 | 2,331,387 | 0.22 % |
| Satellites: | 208 | 145,391 | 0.01 % |
| Simple repeats: | 32,409 | 1,932,561 | 0.18 % |
| Low complexity: | 3,676 | 235,728 | 0.02 % |

**Suppl Table S2. Comparison of genomes of the Rhodophyta**. The genomes published on NCBI (https://www.ncbi.nlm.nih.gov/, accessed on 04/07/2024) and in [1] are reported. *P. palmata* has the third largest genome, the highest number of genes and the highest portion of repeat elements in the Rhodophyta.

| Species | Class/Order | Mu lt | Ext re m | Size (Mb) | GC% | Nr | uni t | N50 (Mb) | Repeats % | BUSCO compl. | PC genes | Year | Ref./NCBI ID |
|---|---|---|---|---|---|---|---|---|---|---|---|---|---|
| *Acrochaetium catenulatum* | F, Acrochaetiales | T | F | 145.72 | | | | | | 65.5 | | 2024 | [1] |
| *Acrochaetium collopodum* | F, Acrochaetiales | T | F | 111.82 | | | | | | 48.2 | | 2024 | [1] |
| *Agarophyton chilense* | F, Gracilariales | T | F | 76.00 | 48.9 | 138 | CN | 1.56 | 66.2 | 75.3 | 7943 | 2023 | [2] |
| *Amphiroa fragilissima* | F, Corallinales | T | F | 146.40 | 42.0 | 2871 | CN | 0.08 | | | | 2024 | ASM4043770v1 |
| *Asparagopsis taxiformis* | F, Bonnemaisoniales | T | F | 142.40 | 49.5 | 3304 | CN | 0.55 | | | 10867 | 2023 | [3] |
| *Audouinella boryana* | F, Acrochaetiales | T | F | 230.40 | | | | | | 69.0 | | 2024 | [1] |
| *Bangia atropurpurea* | B, Bangiales | T | F | 75.17 | | | | | | 73.3 | | 2024 | [1] |
| *Bonnemaisonia californica* | F, Bonnemaisoniales | T | F | 247.60 | 44.0 | 484346 | SC | | | | | 2023 | ASM3235548v1 |
| *Bostrychia calliptera* | F, Ceramiales | T | F | 573.13 | | | | | | 38.4 | | 2024 | [1] |
| *Bostrychia flagellifera* | F, Ceramiales | T | F | 1232.20 | | | | | | 73.7 | | 2024 | [1] |
| *Bostrychia moritziana* | F, Ceramiales | T | F | 588.97 | | | | | | 75.7 | | 2024 | [1] |
| *Bostrychia radicans* | F, Ceramiales | T | F | 1459.82 | | | | | | 81.6 | | 2024 | [1] |
| *Callophyllis flabellulata* | F, Gigartinales | T | F | 88.06 | 45.5 | 97989 | SC | | | | | 2023 | ASM3227052v1 |
| *Caloglossa monosticha* | F, Ceramiales | T | F | 428.32 | | | | | | 76.7 | | 2024 | [1] |
| *Caloglossa ogasawaraensis* | F, Ceramiales | T | F | 443.95 | | | | | | 78.8 | | 2024 | [1] |
| *Caloglossa leprieurii* | F, Ceramiales | T | F | 354.88 | | | | | | 88.6 | | 2024 | [1] |
| *Caloglossa rotundata* | F, Ceramiales | T | F | 154.84 | | | | | | 69.4 | | 2024 | [1] |
| *Caloglossa vieillardii* | F, Ceramiales | T | F | 694.72 | | | | | | 84.3 | | 2024 | [1] |
| *Catenella caespitosa* | F, Gigartinales | T | F | 240.99 | | | | | | 80.0 | | 2024 | [1] |
| *Catenella fusiformis* | F, Gigartinales | T | F | 307.08 | | | | | | 85.1 | | 2024 | [1] |
| *Catenella nipae* | F, Gigartinales | T | F | 233.52 | | | | | | 74.9 | | 2024 | [1] |
| *Chondracanthus exasperatus* | F, Gigartinales | T | F | 32.70 | 41.5 | 117583 | SC | | | | | 2023 | ASM3235821v1 |
| *Chondria armata* | F, Ceramiales | T | F | 507.58 | 45.3 | 2990 | SC | 0.64 | 45.5 | 69.0 | | 2022 | [4] |
| *Chondria dasyphylla* | F, Ceramiales | T | F | 271.92 | | | | | | 75.3 | | 2024 | [1] |
| *Chondrus crispus* | F, Gigartinales | T | F | 104.80 | 52.5 | 925 | SC | 0.24 | 59.0 | 79.9 | 9603 | 2013 | [5] |

| **Species** | **Class/Order** | **Mult** | **Extrem** | **Size (Mb)** | **GC%** | **Nr** | **unit** | **N50 (Mb)** | **Repeats %** | **BUSCO compl.** | **PC genes** | **Year** | **Ref./NCBI ID** |
|---|---|---|---|---|---|---|---|---|---|---|---|---|---|
| *Chromastrum kylinoides* | F, Acrochaetiales | T | F | 92.12 | | | | | | 51.8 | | 2024 | [1] |
| *Chroothece richterianum* | S, Stylonematales | F | F | 75.24 | | | | | | 72.6 | | 2024 | [1] |
| *Cumathamnion decipiens* | F, Ceramiales | T | F | 110.50 | 66.5 | 136873 | SC | | | | | 2023 | ASM3235642v1 |
| *Cyanidiococcus yangmingshanensis* | B, Cyanidiales | F | T | 12.00 | 54.6 | 20 | CH | | | 96.7 | 4832 | 2023 | [6] |
| *Cyanidioschyzon merolae* | B,Cyanidiales | F | T | 16.55 | 55.0 | 20 | CH | 0.86 | 20.0 | 80.6 | 4803 | 2007 | [7] |
| *Cyanidium caldarium* | B,Cyanidiales | F | T | 8.79 | 65.7 | 20 | CH | | | 94.7 | 4870 | 2023 | [6] |
| *Devaleraea mollis* | F, Palmariales | T | F | 775.2 | | 1575647 | SC | | | | | 2023 | ASM3236126v1 |
| *Digenea simplex* | F, Rhodomelaceae | T | F | 299.30 | 43.0 | 8246 | CN | 0.58 | | | | 2019 | [8] |
| *Endocladia muricata* | F, Gigartinales | T | F | 203.90 | 49.5 | 130027 | SC | | | | | 2023 | ASM3235624v1 |
| *Erythrotrichia carnea* | C, Erythropeltales | T | F | 128.01 | | | | | | 56.9 | | 2024 | [1] |
| *Galdieria partita* | B, Galdierales | F | T | 17.80 | 37.5 | 80 | CN | 0.23 | | | 7832 | 2022 | [9] |
| *Galdieria phlegrea* | B, Galdierales | F | T | 14.87 | 37.5 | 108 | CN | 0.20 | | 95.1 | 6125 | 2019 | [10] |
| *Galdieria sulphularia* | B, Galdierales | F | T | 14.50 | 40.2 | 76 | SC | 0.19 | | 95.7 | 7021 | 2023 | [6] |
| *Gracilaria caudata* | F, Gracilariales | T | F | 30.28 | 49.9 | 5535 | CN | 0.21 | 45.7 | 73.0 | 8737 | 2023 | [2] |
| *Gracilaria changii* | F, Gracilariales | T | F | 35.80 | 50.6 | 10853 | CN | | | 70.0 | 10912 | 2018 | [11] |
| *Gracilaria domingensis* | F, Gracilariales | T | F | 77.70 | 49.7 | 684 | SC | 0.19 | | 87.5 | 11437 | 2022 | [12] |
| *Gracilaria gracilis* | F, Gracilariales | T | F | 72.49 | 46.6 | 279 | CN | 0.56 | 60.7 | 77.3 | 9460 | 2023 | [2] |
| *Gracilaria vermiculophylla* | F, Gracilariales | T | F | 44.95 | 49.5 | 4240 | CN | 2.56 | 48.3 | 65.1 | 6807 | 2023 | [13], [2] |
| *Gracilariopsis chorda* | F, Gracilariales | T | F | 92.18 | 49.3 | 1211 | CN | 0.22 | 61.0 | 87.6 | 10806 | 2018 | [14] |
| *Gracilariopsis lemaneiformis* | F, Gracilariales | T | F | 88.98 | 48.1 | 13825 | SC | 0.03 | | | 9281 | 2018 | [15] |
| *Hildenbrandia prototypus* | F, Hildenbrandiales | T | F | 145.56 | | | | | | 79.2 | | 2024 | [1] |
| *Hymenocladiopsis crustigena* | F, Rhodymeniales | T | T | 134.98 | | | | | | 76.9 | | 2024 | [1] |
| *Hypoglossum anomalum* | F, Ceramiales | T | T | 666.44 | | | | | | 67.1 | | 2024 | [1] |
| *Jania rubens* | F, Corallinales | T | F | 251.43 | | | | | | 80.8 | | 2024 | [1] |
| *Kappaphycus alvarezii* | F, Gigartinales | T | F | 336.1 | 45.5 | 888 | CN | 0.85 | | | | 2020 | ASM220596v3 |
| *Kylinia rosulata* | F, Acrochaetiales | T | F | 137.20 | | | | | | 71.8 | | 2024 | [1] |
| *Mastocarpus papillatus* | F, Gigartinales | T | F | 31.4 | 53.5 | 33839 | SC | | | | | 2023 | ASM3235614v1 |

| Species | Class/Order | Mult | Extrem | Size (Mb) | GC% | Nr | unit | N50 (Mb) | Repeats % | BUSCO compl. | PC genes | Year | Ref./NCBI ID |
|---|---|---|---|---|---|---|---|---|---|---|---|---|---|
| *Microcladia borealis* | F, Ceramiales | T | F | 832.4 | 43.5 | 604122 | SC | | | | | 2023 | ASM3227414v1 |
| *Myriogramme manginii* | F, Ceramiales | T | T | 347.58 | | | | | | 75.7 | | 2024 | [1] |
| *Nemalionopsis parkeri* | F, Thoreales | T | F | 81.72 | | | | | | 69.8 | | 2024 | [1] |
| *Neoporphyra haitanensis* | B, Bangiales | T | F | 49.67 | 70.1 | 15 | CH | 7.80 | 31.6 | 85.8 | 9496 | 2022 | [16] |
| *Odonthalia washingtoniensis* | F, Ceramiales | T | F | 298.5 | 40.5 | 108059 | SC | | | | | 2023 | ASM3235823v1 |
| *Opuntiella californica* | F, Gigartinales | T | F | 28.9 | 39.5 | 83713 | SC | | | | | 2023 | ASM3227064v1 |
| *Palmaria decipiens* | F, Palmariales | T | T | 349.78 | | | | | | 68.6 | | 2024 | [1] |
| ***Palmaria palmata*** | **F, Palmariales** | **T** | **F** | **1050.31** | **51.9** | **1015** | **CN** | **2.75** | **91.3** | **72.1** | **9641** | | **This study** |
| *Phyllophora antarctica* | F, Gigartinales | T | T | 348.08 | | | | | | 47.1 | | 2024 | [1] |
| *Polyneura hilliae* | F, Ceramiales | T | F | 545.52 | | | | | | 78.4 | | 2024 | [1] |
| *Porolithon onkodes* | F, Corallinales | T | F | 226.2 | 46.0 | 68138 | SC | | | | | 2023 | PU_POon_v1 |
| *Porphyra lucasii* | B, Bangiales | T | F | 275.75 | | | | | | 65.1 | | 2024 | [1] |
| *Porphyra miniata* | B, Bangiales | T | F | 302.89 | | | | | | 90.2 | | 2024 | [1] |
| *Porphyra plocamiestris* | B, Bangiales | T | T | 185.31 | | | | | | 86.3 | | 2024 | [1] |
| *Porphyra pulchella* | B, Bangiales | T | F | 197.48 | | | | | | 82.4 | | 2024 | [1] |
| *Porphyra umbilicalis* | B, Bangiales | T | F | 87.70 | 65.8 | 2125 | SC | 0.13 | 43.9 | 42.4 | 13125 | 2017 | [17] |
| *Porphyridium purpureum* | B, Porphyridiales | F | F | 22.20 | 55.5 | 52 | CN | 1.90 | 3.9 | 88.5 | 8355 | 2019 | [18] |
| *Pterothamnion heteromorphum* | F, Ceramiales | T | F | 281.2 | 45.0 | 272808 | SC | | | | | 2023 | ASM3235829v1 |
| *Pulvinaster venetus* | C, Compsopogonales | T | F | 95.91 | | | | | | 40.8 | | 2024 | [1] |
| *Pyropia haitanensis* | B, Bangiales | T | F | 53.30 | 67.9 | 5 | CH | 5.80 | 24.2 | 85.5 | 10903 | 2020 | [19] |
| *Pyropia yezoensis* | B, Bangiales | T | F | 108.00 | 64.8 | 3 | CH | 0.34 | 48.0 | 84.0 | 12855 | 2020 | [20] |
| *Rhodachlya madagascarensis* | F, Rhodachlyales | T | F | 143.72 | | | | | | 60.8 | | 2024 | [1] |
| *Rhodenigma contortum* | F, Rhodogorgonales | T | F | 183.41 | | | | | | 74.9 | | 2024 | [1] |
| *Rhodochorton purpureum* | F, Acrochaetiales | T | F | 219.13 | | | | | | 60.4 | | 2024 | [1] |
| *Rhodosorus marinus* | S, Stylonematales | F | F | 31.2 | 49.0 | 17 | CN | 3.2 | | | 8392 | 2023 | [21] |
| *Rhodothamniella floridula* | F, Palmariales | T | F | 183.84 | | | | | | 63.5 | | 2024 | [1] |
| *Rhodymenia pacifica* | F, Rhodymeniales | T | F | 67.8 | 40.5 | 142089 | SC | | | | | 2023 | ASM3227032v1 |

| Species | Class/Order | Mult | Extrem | Size (Mb) | GC% | Nr | unit | N50 (Mb) | Repeats % | BUSCO compl. | PC genes | Year | Ref./NCBI ID |
|---|---|---|---|---|---|---|---|---|---|---|---|---|---|
| *Sarcodiotheca gaudichaudii* | F, Gigartinales | T | F | 124.3 | 40.5 | 327224 | SC | | | | | 2023 | ASM3235598v1 |
| *Thorea hispida* | F, Thorales | T | F | 126.61 | | | | | | 77.7 | | 2024 | [1] |
| *Thorea riekei* | F, Thorales | T | F | 111.04 | | | | | | 71.4 | | 2024 | [1] |
| *Thorea violacea* | F, Thorales | T | F | 104.15 | | | | | | 72.2 | | 2024 | [1] |
| *Vertebrata fucoides* | F, Ceramiales | T | F | 500.18 | | | | | | 76.5 | | 2024 | [1] |

Class: B= Bangiophyceae, C= Compsopogonophyceae; F= Florideophyceae, S= Stylonematophyceae; Multicell (T/F): individuals are multicellular (true or false); Extrem (T/F): extremophile (true or false); Nr: number of chromosomes or scaffold or contigues; unit: it is the unit of measurement referred to the previous column, CH= chromosome, CN= contig, SC= scaffold; BUSCO compl.: BUSCO completeness; PC genes: number of protein coding genes; Year: year of publication. Ref/NCBI ID: reference to a paper when available or assembly identifier from NCBI.

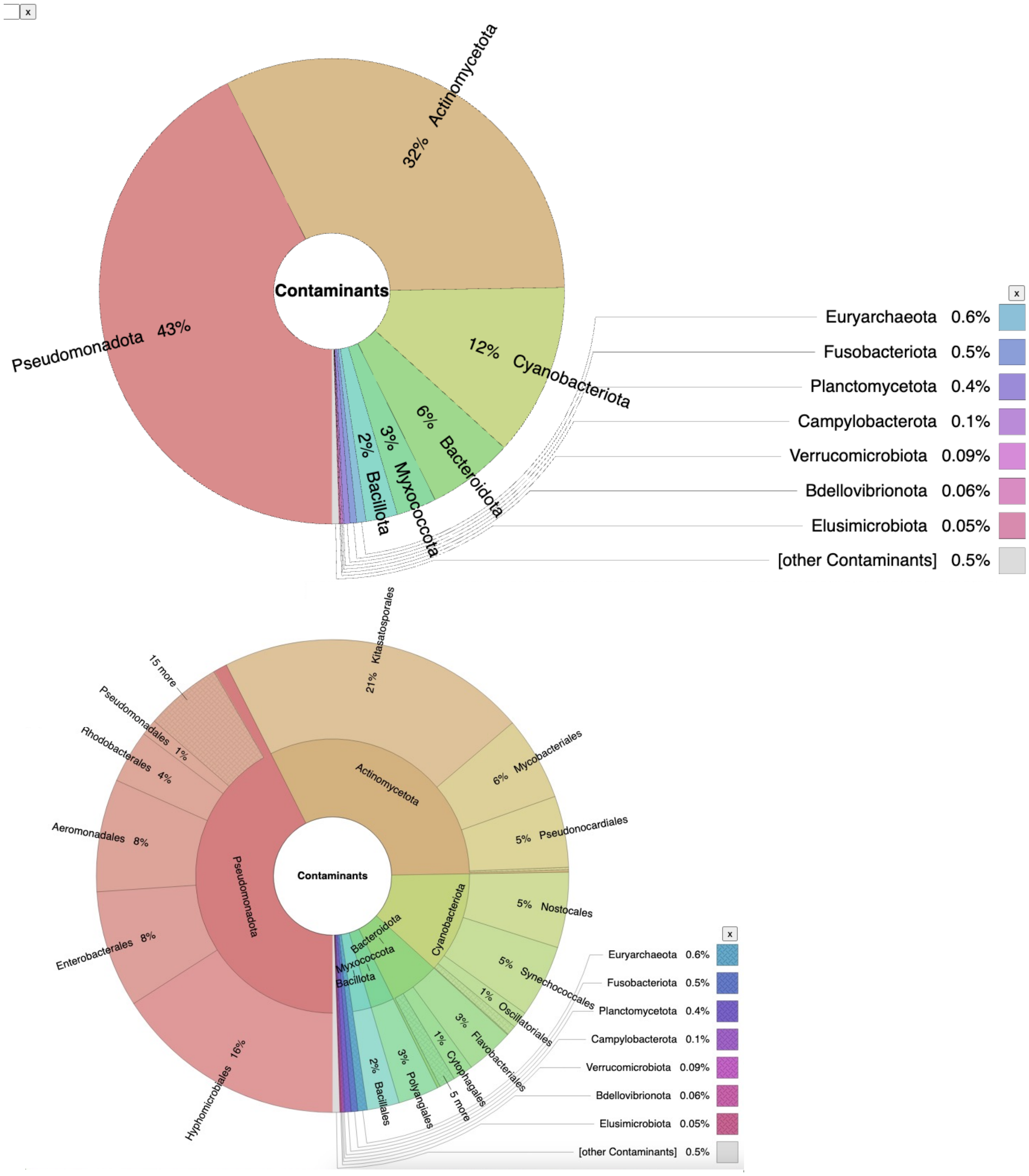


**Supplementary Figure S2. Classification of contaminant DNA** at the level of a. Phylum and b. Order.

**Supplementary Table S3. Number of SNPs per location.**

| Id | Location | Nr. SNPs |
|---|---|---|
| P38 | Fornaes, DNK | 4944 |
| P36 | Fornaes, DNK | 4909 |
| P33 | Fornaes, DNK | 4882 |
| P13 | Famjin, FRO | 4511 |
| P41 | Loddbukta, NOR | 4423 |
| P7 | Famjin, FRO | 4257 |
| P44 | Aursvika, NOR | 4204 |
| P39 | Loddbukta, NOR | 4180 |
| P42 | Loddbukta, NOR | 4156 |
| P10 | Oyragjogv, FRO | 4019 |
| P4 | Oyragjogv, FRO | 4012 |
| P11 | Oyragjogv, FRO | 3941 |
| P14 | Famjin, FRO | 3855 |
| P5 | Oyragjogv, FRO | 3846 |
| P17 | Roundstone, IRL | 3744 |
| P31 | Ballycastle, NIR | 3723 |
| P30 | Ballycastle, NIR | 3700 |
| P15 | Roundstone, IRL | 3699 |
| P12 | Famjin, FRO | 3485 |
| P32 | Ballycastle, NIR | 3301 |
| P9 | Hvitanes, FRO | 3251 |
| P8 | Hvitanes, FRO | 3242 |
| P16 | Roundstone, IRL | 3200 |
| P6 | Famjin, FRO | 3191 |
| P25 | Saint-Brieuc, FRA | 3189 |
| P2 | Hvitanes, FRO | 2911 |
| P28 | Saint-Brieuc, FRA | 2835 |
| P27 | Saint-Brieuc, FRA | 2579 |
| P3 | Hvitanes, FRO | 2522 |
| P18 | Roundstone, IRL | 2086 |
| P20 | Roundstone, IRL | 2039 |
| P19 | Roundstone, IRL | 1443 |
| P1 | Ballyvaughan, IRL | 924 |

**Supplementary Table S4. List of unique SNPs per locations.** The first part of the name SNP is the contig, followed by the position, the reference and the alternate allele value.

| | |
|---|---|
| Hvítanes, FRO | ptg000014l_3484421_A_T |
| Fornæs, DNK | ptg000008l_1044130_A_T |
| | ptg000008l_1044133_A_G |
| | ptg000008l_1044155_G_T |
| | ptg000008l_1044263_T_A |
| | ptg000008l_1044347_A_T |
| | ptg000008l_1044675_C_T |
| | ptg000014l_633892_C_T |
| | ptg000020l_1523690_A_G |
| | ptg000020l_1524726_G_C |
| | ptg000020l_4406100_C_T |
| | ptg000023l_838977_G_A |
| | ptg000023l_1696789_G_A |
| | ptg000023l_2874010_A_T |
| | ptg000023l_2900002_C_T |
| | ptg000023l_2903651_A_G |
| | ptg000023l_2903676_A_G |
| Ballycastle, NIR | ptg000014l_3484601_T_C |
| | ptg000023l_2877255_G_T |
| | ptg000023l_2877423_A_C |
| Saint-Brieuc, FRA | ptg000006l_462698_A_T |
| | ptg000016l_57733_T_C |
| | ptg000021l_2728099_C_A |
| | ptg000021l_2728180_A_C |
| | ptg000022l_585379_C_T |
| | ptg000023l_2897747_T_C |
| | ptg000023l_2898794_C_T |
| | ptg000023l_2899710_G_C |
| | ptg000023l_2899788_C_A |
| | ptg000023l_2899833_C_T |
| | ptg000023l_2900119_T_A |
| | ptg000023l_2900778_G_C |
| | ptg000023l_3148107_G_C |
| | ptg000023l_3148215_A_T |

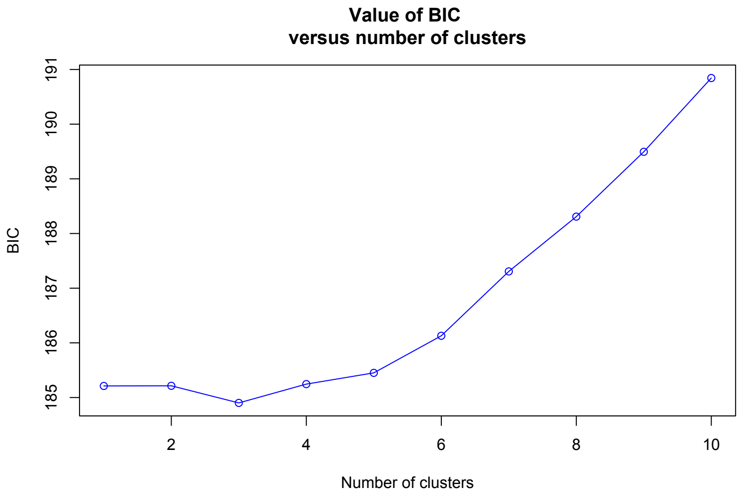


**Supplementary Figure S3. BIC against the number of clusters.** Discriminant analysis of principal components (DAPC) using 9,885 SNPs of 33 genotypes was performed for 1–33 clusters and the Bayesian information criterion (BIC) was used to identify the optimal number of clusters/ancestral populations to describe the population: k= 3.
